# The order of trait emergence in the evolution of cyanobacterial multicellularity

**DOI:** 10.1101/570788

**Authors:** Katrin Hammerschmidt, Giddy Landan, Fernando Domingues Kümmel Tria, Jaime Alcorta, Tal Dagan

## Abstract

The transition from unicellular to multicellular organisms is one of the most significant events in the history of life. Key to this process is the emergence of Darwinian individuality at the higher level: groups must become single entities capable of reproduction for selection to shape their evolution. Evolutionary transitions in individuality are characterized by cooperation between the lower level entities and by division of labor. Theory suggests that division of labor may drive the transition to multicellularity by eliminating the trade-off between two incompatible processes that cannot be performed simultaneously in one cell. Here we examine the evolution of the most ancient multicellular transition known today, that of cyanobacteria, where we reconstruct the sequence of ecological and phenotypic trait evolution. Our results show that the prime driver of multicellularity in cyanobacteria was the expansion in metabolic capacity offered by nitrogen fixation, which was accompanied by the emergence of the filamentous morphology and succeeded by a reproductive life cycle. This was followed by the progression of multicellularity into higher complexity in the form of differentiated cells and patterned multicellularity.

**Significance Statement:** The emergence of multicellularity is a major evolutionary transition. The oldest transition, that of cyanobacteria, happened more than 3 to 3.5 billion years ago. We find N_2_ fixation to be the prime driver of multicellularity in cyanobacteria. This innovation faced the challenge of incompatible metabolic processes since the N_2_ fixing enzyme (nitrogenase) is sensitive to oxygen, which is abundantly found in cyanobacteria cells performing photosynthesis. At the same time, N_2_-fixation conferred an adaptive benefit to the filamentous morphology as cells could divide their labour into performing either N_2_-fixation or photosynthesis. This was followed by the culmination of complex multicellularity in the form of differentiated cells and patterned multicellularity.

## Introduction

Multicellularity is considered a characteristic trait of eukaryotes, but has evolved independently several times in diverse prokaryote taxa, including actinobacteria, myxobacteria, and cyanobacteria (Bonner 1998). Bacterial multicellularity ranges from transient associations, such as colonies, biofilms and cellular aggregations, to permanent multicellular forms (Shapiro 1988)

Instances of multicellular bacterial species present the major traits of eukaryotic multicellularity, including cell-to-cell adhesion, peri- or cytoplasmic continuity, intercellular communication, patterning, programmed cell death (PCD), and division of labor (Claessen et al. 2014). Aggregative forms of multicellularity are common among bacterial species, for example, those that form a biofilm under specific external conditions (Tarnita et al. 2013). *Bacillus subtilis*, for instance, forms biofilms upon nutrient deprivation in which cells differentiate into motile, matrix producing, or spore cells depending on the environmental cues (Claessen et al. 2014). Notably, cell differentiation in aggregates is adaptive at the level of the individual cell as it directly confers a fitness benefit to that particular cell. In contrast, under true division of labor, cells are interdependent upon each other and specialize in performing complementary tasks. These tasks, e.g., somatic functions or PCD, are not beneficial on the level of the individual cell, but are advantageous for the colony; thus, they are emergent properties on a higher level of organization (van Gestel et al. 2015).

True division of labor in bacteria is best described in actinobacteria and cyanobacteria (van Gestel et al. 2015). In cyanobacteria, the most complex of the filamentous species can differentiate up to five different cell types: vegetative (photosynthetic) cells, akinetes (spore-like cells), hormogonia (reproductive, motile filaments), necridia (dead cells resulting from PCD/apoptosis for hormogonia release), and heterocysts (Claessen et al. 2014; Herrero et al. 2016). Heterocysts differentiate under nitrogen deprivation and are specialized in nitrogen (N_2_) fixation by the enzyme nitrogenase (Frías et al. 1994). As this enzyme is sensitive to oxygen (O_2_), these cells are characterized by the absence of oxygenic photosynthesis and by a thick cell wall, which maintains an anaerobic environment. Heterocysts and vegetative cells in the filament are metabolically interdependent with the heterocysts providing combined nitrogen to the other cells within the filament and receiving fixed carbon compounds in return. Heterocysts cannot reproduce hence they represent a prime example for emergent traits on the level of a multicellular organism.

Cyanobacteria possess the hallmark traits reminiscent of complex eukaryotic multicellularity, making the order of trait emergence essential for understanding the origin of higher-level complexity in organismal evolution. Here we infer the evolutionary trajectory of the emergence of traits in the evolution of multicellularity in cyanobacteria.

## Materials and Methods

### Data

The primary data underlying this study consists of the genomic sequences and phenotypic traits of 199 representative cyanobacterial species. These were selected from the available genomes so that the number of represented taxa will be as large as possible and genus-level redundancy will be reduced (see supplementary table S1 for the complete list of species).

### Phenotypic traits

Phenotypic traits were chosen for their potential relevance to the evolution of multicellularity in cyanobacteria, such as environmental factors that might facilitate multicellularity and markers that are indicative for the transition to multicellularity (table 1).

**Table 1.**
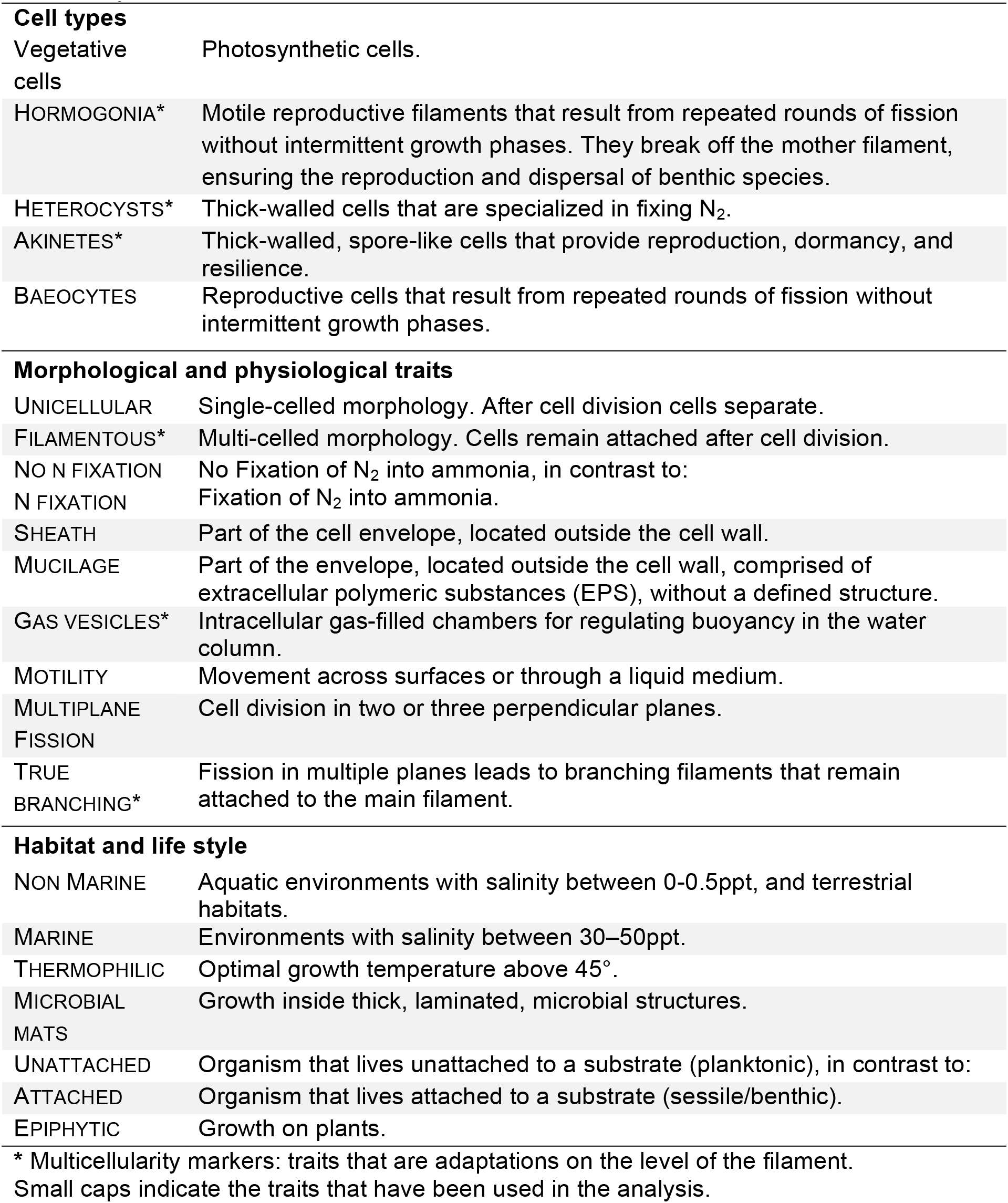
Description of cyanobacterial cell types, morphological and physiological traits, their habitat and life style.

Information on presence and absence of traits was obtained from the published literature and from the Pasteur Culture Collection of cyanobacteria, extending the work by Uyeda et al. 2016, and coded as binary trait states. Traits included morphology (unicellular, filamentous), nitrogen fixation (no N_2_ fixation, N_2_ fixation), habitat (marine/ non marine), baeocytes, hormogonia, thermophilic, akinetes, heterocysts, true branching, epiphytic, microbial mats, attached/ unattached, sheath, mucilage, gas vesicles, motility, and multiplane fission (table 1, supplementary table S1).

### Protein families and alignments

The cyanobacteria protein families were constructed from completely sequenced genomes available in RefSeq database (O’Leary et al. 2016; ver. May 2016). For the construction of protein families, at the first stage, all protein sequences annotated in the genomes were blasted all-against-all using stand-alone BLAST (Altschul et al. 1990) ver. 2.2.26. Protein sequence pairs that were found as reciprocal best BLAST hits (rBBHs) (Tatusov et al. 1997) with a threshold of E-value ≤ 1×10^−5^ were further compared by global alignment using needle (Rice et al. 2000). Sequence pairs having ≥30% identical amino acids were clustered into protein families using the Markov clustering algorithm (MCL) (Enright et al. 2002) ver. 12-135 with the default parameters. Multiple-copy gene families were discarded, resulting in an initial dataset of 18,873 single-copy gene families.

Gene families were then extended to include homologous sequences from non-cyanobacteria species, serving as outgroups for rooting purposes. We identified outgroup homologues by an rBBH analysis of the *Scytonema hofmanni* PCC 7110 genome (the most widely present species in the initial gene family dataset) against 26 high quality non-cyanobacteria genomes: Vampirovibrionia (12 genomes) and Sericytochromatia (2) (Soo et al. 2017; Carnevali et al. 2019), the closest phyla Margulisbacteria (6), Saganbacteria (2), Fusobacteria (1) and Firmicutes (1) (according to (Carnevali et al. 2019) and (Zhu et al. 2019); one reference anoxygenic photosynthetic genome from *Chloroflexus aurantiacus* J-10-fl and the *Escherichia coli* str. K-12 substr. MG1655 genome (see supplementary table S1). The number of gene families with homologs ranged between 204 and 451 for the 26 outgroup genomes. We selected six of these outgroups for further analyses: *Vampirovibrio chlorellavorus, Chloroflexales, Obscuri-PALSA-1081, Sericytochromatia-UBA7694, Bacillus subtilis,* and *Margulis-GWF2-35-9.* Protein sequences of these families were aligned using MAFFT version 7.027b employing the L-INS-i strategy (Katoh & Standley 2013). The alignments are available in supplementary material online.

### Species tree reconstruction

The sequence data for the reconstruction of the cyanobacterial species tree consisted of 14 single-copy gene families that are present in all 199 cyanobacteria genomes and any of the six outgroup genomes. The species tree was inferred using IQ-TREE (Nguyen et al. 2015) in a partitioned analysis over the concatenated alignment of the 14 gene alignments (iqtree version 1.6.6.b with parameters -t BIONJ -keep-ident -mset LG -madd LG4X–spp). The unrooted species tree was rooted on the branch leading to the outgroup. The species tree is available in supplementary material online.

### Gene trees reconstruction

To evaluate the robustness of inferences drawn from the species tree, we also reconstructed gene trees to provide a large sample of comparisons to the species tree. The gene trees dataset consisted of 553 single-copy gene families present in at least one genome from both sides of the root of the species tree, and at least one of the six outgroup species. Gene trees were inferred using IQ-TREE (Nguyen et al. 2015) version 1.6.6.b with parameters -t BIONJ - keep-ident -mset LG -madd LG4X). Trees were rooted on the branch separating the outgroup from the ingroup. A total of 138 trees where the outgroup sequences did not form a single partition were discarded. The gene trees are available in supplementary material online.

### Inference of trait order

The traits presence/absence pattern was mapped onto the rooted species tree. Most of the traits in our study are rather complex (i.e., they involved multiple genes), hence their emergence in the evolution of cyanobacteria is expected to be a rare event. Accordingly, we used a parsimonious reconstruction approach and assigned the origin of a trait to the most recent species tree node where the trait is present in any of the node’s descendants. Consistency index (CI) and retention index (RI) for each trait were calculated using the PHYLIP program pars (Felsenstein 2005) as described in Farris (1989) The species tree was traversed from root to tips to determine the order of trait emergence. For each pair of traits we tested whether the order observed in the species tree was reproduced in gene trees with at least two species displaying each of the two traits. For that purpose, we repeated the trait-order analysis with the set of single-copy rooted gene trees, including gene families that do not span the full taxa set. The agreement of the gene trees with the conclusion based on the species trees is calculated as the proportion of gene trees where the trait order is the same as in the species tree.

## Results and Discussion

To reconstruct the order of trait emergence in the evolution of cyanobacterial multicellularity, we evaluated 21 phenotypic traits variably present in 199 cyanobacterial species (table 1, supplementary table S1). The ability to perform photosynthesis is not included in our study as it is universal to all cyanobacteria hence it is considered an ancestral trait (Garcia Pichel et al. 2020). We inferred a species tree from a partitioned analysis of 14 single copy core genes that are present in all 199 species. The species tree was rooted by inclusion of outgroup sequences from six bacterial species (see methods), with the root separating the genus *Gloeobacter* from all other cyanobacteria.

The rooting of cyanobacteria is a thorny issue, with two competing clades put forward as basal lineages. Phylogenetic studies based on single gene phylogenies indicated *Gloeobacter* to be a basal lineage within cyanobacteria (e.g., Bhattacharya & Medlin 1995; Honda et al. 1999). The *Gloeobacter* basal position is in agreement with the cellular characteristics of species in this genus, which are lacking well-defined thylakoids (Rippka et al. 1974; Mareš et al. 2019) that are considered a derived structure in the phylum. Indeed, several studies in the literature thus used *Gloeobacter* for rooting the cyanobacterial species tree (e.g., Shih et al. 2012; Shi & Falkowski 2008; Dagan et al. 2013; Sánchez-Baracaldo et al. 2014). Nonetheless, other studies, which used midpoint rooting or minimal ancestor deviation (MAD) to root the cyanobacteria species tree, position the root on the branch separating between the pico-cyanobacteria (*Synechococcus* & *Prochlorococcus,* and hereafter SynPro clade) and the remaining species (Szöllősi et al. 2012; Tria et al. 2017). This branch, however, is typically long in gene trees, as well as the species trees, hence it may reflect a Long Branch Attraction (LBA) artifact (defined in Felsenstein 1978).

To evaluate whether the rooting placement is robust, we conducted gene-tree support analyses by reconstructing trees for 553 cyanobacterial single-copy gene families along with homologs from six different outgroup species (fig. 1, and methods). We then extracted the rooted cyanobacterial subtree, while discarding 138 trees where the outgroup species formed more than one group. We next characterize the trees by the pattern of the three cyanobacterial subgroups: *Gloeobacter* (2 spp.), SynPro clade (32 spp.), and Other Cyanobacteria (165 spp.), discarding 142 genes that are present in less than two genomes for each group. First, we considered gene trees in which no group is present on both sides of the root. The group appearing on its own as a lineage originating at the root is considered a basal group and the tree is supporting a root located on the branch separating it from the others (left column in fig. 1). Next, if only one group appears on both sides of the root, this group is labeled as ancestral, and supports a root position within it (right column in fig. 1). We found no gene trees where more than one group appears on both sides of the root. We note that the gene trees may be discordant with the species tree, and that any of the groups may seem to be paraphyletic, either due to methodological artifacts (e.g. LBA involving the SynPro clade) or due to biological processes such as lateral gene transfer (LGT).

**Figure 1.**
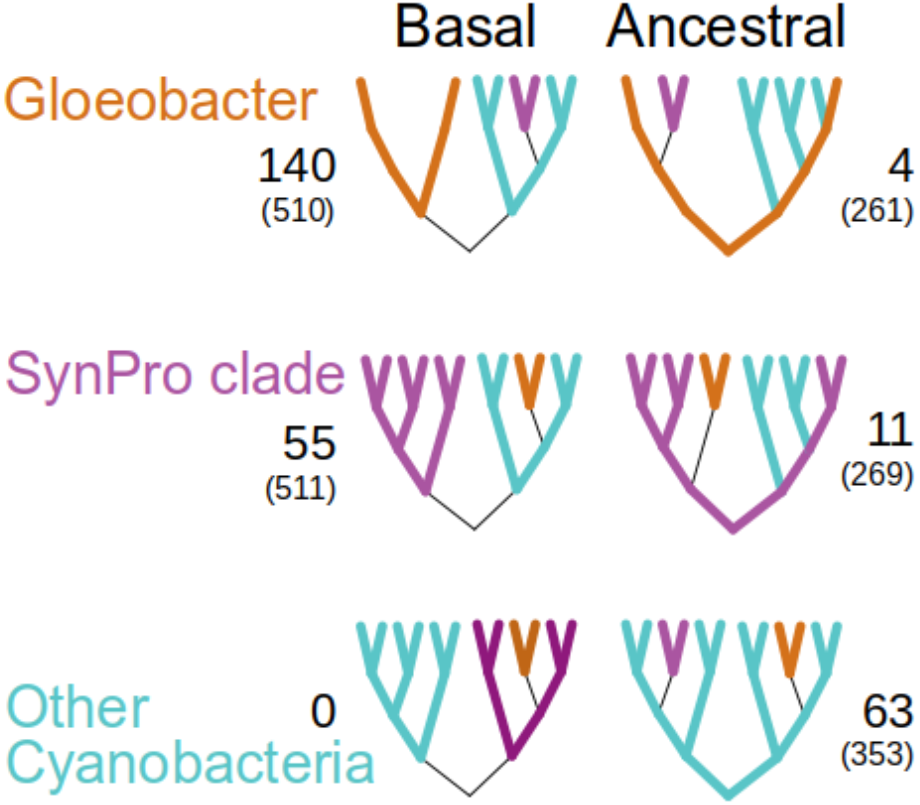
Support for three possible basal/ancestral cyanobacterial groups in 273 rooted gene trees. The number of gene families supporting each type of rooted topology is given (median alignment length is shown in parenthesis).

Our results reveal that the majority of gene trees identify the *Gloeobacter* lineage as a distinct basal lineage stemming from the root, thus supporting the outgroup rooting of the species tree on that branch.

### The order of trait emergence in cyanobacterial evolution

We infer the order of trait emergence in cyanobacterial evolution by mapping the traits onto the rooted species tree. The *origin node* corresponding to each trait was assigned as the most recent node where the trait is present in any of the node’s descendants. This is a conservative approach in that for each trait it allows for a single origin with possible subsequent losses (Dagan & Martin 2007). Our approach furthermore assumes that the traits in our analysis are vertically inherited rather than acquired by lateral transfer. Indeed, previous phylogenomic studies indicated that gene acquisition by lateral gene transfer is frequent in the evolution cyanobacteria (Zhaxybayeva et al. 2009; Dagan et al. 2013) The traits we included in our analysis, however, are complex phenotypic traits that are the product of multiple genes whose expression is well coordinated with other physiological processes in the cell. Such traits are unlikely to be acquired by lateral gene transfer, because their retention requires that multiple coding and regulation elements will be functional on arrival (Cohen et al. 2011). Nonetheless, since complex systems may have been sporadically and rarely transferred (e.g., Brinkmann et al. 2018), we quantified the level of homoplasy and synapomorphy in the evolution of the studied traits by the consistency and retention indices (Farris 1989). Our results for most of the traits revealed high RI values (median 0.61) and low CI values (median 0.048) (supplementary table S1). This pattern is as expected for complex traits whose evolution is best characterized by a single origin and differential loses.

We next compare the origin nodes for pairs of traits. When the origin of two traits is assigned to the same node, we label the two traits as ‘simultaneous’ at the resolution of the current taxa sample. When an origin node of a trait is a descendent of a second trait’s origin node, we conclude that the first trait emerged earlier. A third possibility is that the origin nodes of two traits are not nested, but this relationship was not observed in any of the 210 trait pairs. The order of trait emergence is depicted in fig. 2.

**Figure 2.**
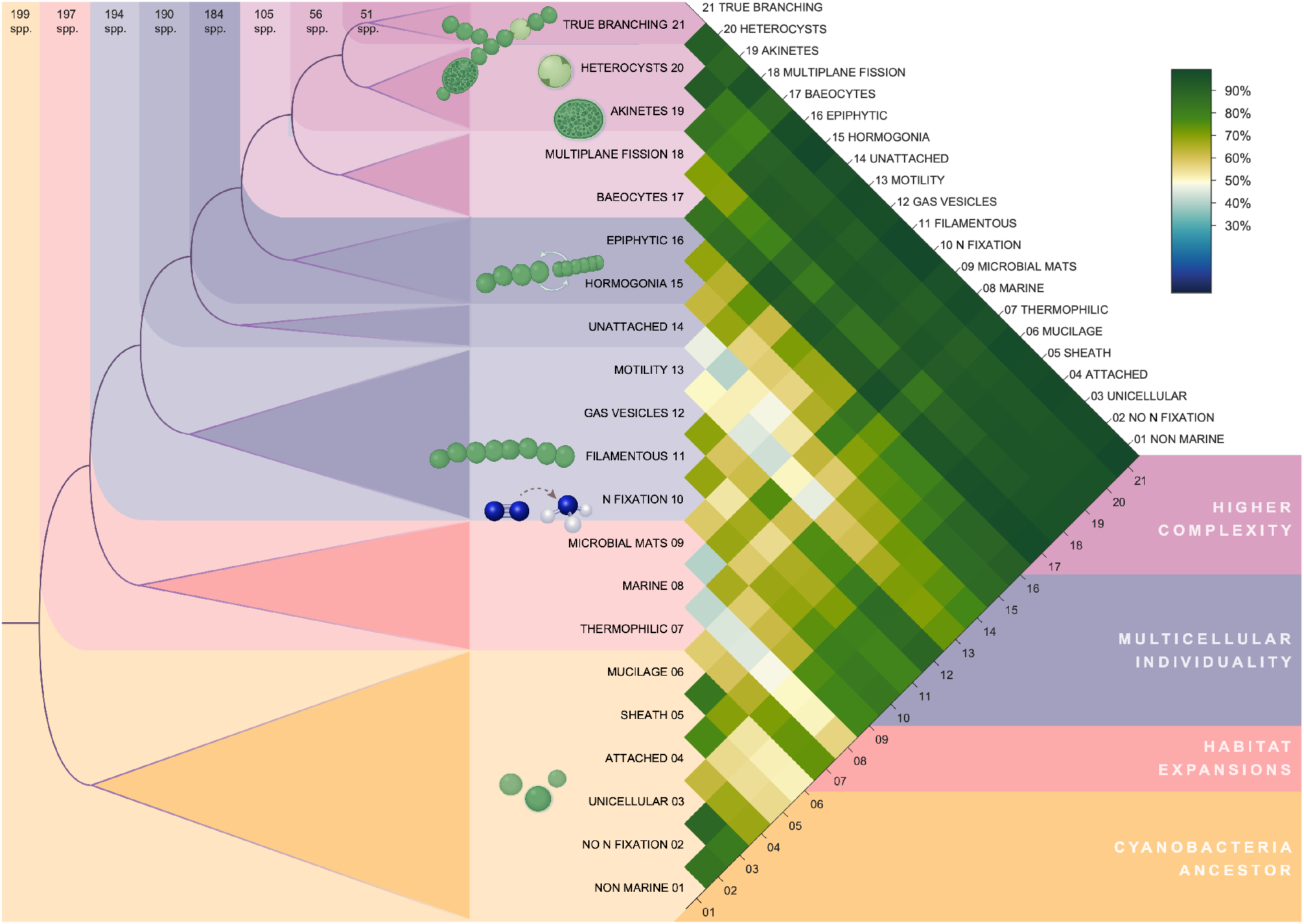
Order of trait emergence. Left: traits and their inferred origin node from the rooted species tree. Colors mark traits with a common origin node (note that the order of traits within the colored blocks is arbitrary). Colored boxes are nested, i.e., earlier traits are present also in the nested colors. Right: Frequency of gene trees in agreement with the relative order of pairs of traits. Cells in the matrix are shaded according to the color bar on the right.

The order of trait emergence is inferred from a species tree, yet, phylogenetic inference of species phylogenies based on the concatenation of only few core genes may suffer from a low resolution of the phylogenetic signal (e.g., Dagan & Martin 2006; Thiergart et al. 2014). To evaluate the robustness of the trait order derived from the species tree, we repeated the analysis by testing the trait order inference in individual gene trees. For that purpose, we considered the set of single-copy gene families where the gene is present in at least one species from both sides of the species tree root and at least one outgroup species. The gene trees were rooted by the outgroup, and the order of pairs of traits determined. The analysis of a large sample of gene trees provides a statistical view not possible in a single species tree, but potentially introduces contradicting inferences due to both biological reasons (LGT) and methodological uncertainties (alignment quality and phylogenetic artifacts such as LBA). In fig. 2 we report the percentage of gene trees that reproduces the species tree ordering. The vast majority of trait pair orderings are observed also in more than 50% of the individual gene trees. Excluding the sequences from the SynPro clade from the gene trees led to slightly better agreement of the gene trees with the order observed in the species tree (supplementary fig. S1), suggesting that LBA artifacts associated with the SynPro clade may lead to disagreement between the gene and species trees. We note that gene trees that are discordant with the species tree generally had shorter alignment length and lower bootstrap support (supplementary tables S2 and S3). Our results thus show a high level of agreement between the species tree and the gene trees, as is expected under an overall homogeneous yet low frequency of LGT during bacterial evolution (Dagan & Martin 2007).

In what follows we divide the inferred order of trait emergence into four temporal phases: (phase i) the cyanobacteria ancestor (traits 1-6); (phase ii) habitat expansions (traits 7-9); (phase iii) the transition to multicellular individuality (traits 10-16); and (phase iv) the evolution of higher complexity (traits 17-21).

### The cyanobacterial ancestor and subsequent habitat expansion

The rooted tree topology supports the view that the cyanobacterial ancestor was characterized by traits that include UNICELLULAR and NO N-FIXATION (fig. 2). The ancestral state of both traits as preceding the emergence of filamentous forms is debated in the literature - whereas one study suggested the ancestor to be unicellular and the filamentous morphology to arise in independent lineages of the cyanobacterial tree (Sánchez-Baracaldo et al. 2005), another view posed that the filamentous morphology evolved early during cyanobacterial evolution and was subsequently lost and regained several times (Schirrmeister et al. 2011). There are also claims that the last cyanobacterial common ancestor already fixed N_2_ (Tomitani et al. 2006), there are however others that concluded that it could not fix N_2_ and that cyanobacteria must have acquired this trait several times independently (Sánchez-Baracaldo et al. 2005).

We can further deduce from our data that the cyanobacterial ancestor lived ATTACHED and possessed a SHEATH and MUCILAGE. Whether the cyanobacteria ancestor lived ATTACHED is a matter of debate and opposing views on the topic have been published (Sánchez-Baracaldo et al. 2005; Uyeda et al. 2016; Schopf 1993; Garcia-Pichel 1998; Sánchez-Baracaldo 2015)

SHEATH and MUCILAGE are forms of extracellular polymeric substances (EPS), located outside the cell wall, which in today’s cyanobacteria are mainly involved in protecting the cell from various stresses, such as UV and desiccation (Ehling-Schulz & Scherer 1999; Potts 1994). Furthermore, we find the cyanobacteria ancestor to have most likely inhabited a NON MARINE environment, agreeing with studies that suggest that early cyanobacteria lived in freshwater or terrestrial habitats and subsequently diverged into marine environments (Dagan et al. 2013; Uyeda et al. 2016).

The second phase in cyanobacterial evolution is the expansion of the cyanobacterial habitat, indicated by the traits MARINE and THERMOPHILIC, which are both inferred to simultaneously occur with the ability to form MICROBIAL MATS. MICROBIAL MATS are dense communities (Stal 1995) that typically present a laminated segregation of functional types. They are often formed by cyanobacteria and are frequently found in extreme habitats, such as deserts or hot springs, characterized by temperatures between 30°C to 73°C (e.g., (Cox et al. 2011)).

### The emergence of N_2_ fixation is at the origin of cyanobacterial multicellularity

Phase iii in our reconstruction comprises three sets of cyanobacterial traits (fig. 2). First, the simultaneous emergence of the FILAMENTOUS morphology, N FIXATION, GAS VESICLES and MOTILITY, followed by the trait UNATTACHED, and lastly by the co-occurrence of HORMOGONIA and EPIPHYTIC. During cyanobacterial N_2_ fixation, molecular dinitrogen (N_2_) is reduced to ammonia (NH_3_), a process that is catalyzed by the enzyme nitrogenase. Whereas present day cyanobacteria, other microorganisms, and most plants are able to take up nitrogen in various combined forms, such as nitrate, ammonium, organic nitrogen, or urea, these combined forms of nitrogen are scarce in most environments (e.g., open oceans or terrestrial habitats (Zehr 2011)). Combined nitrogen, which is critical for the biosynthesis of amino and nucleic acids, was likely a limiting resource in the early Earth environment (Kasting & Siefert 2001).

The realization of the full metabolic potential of N_2_ fixation, however, faced the challenge of the incompatibility of nitrogenase with intracellular oxygen (Gallon 1981). When the cyanobacterial ancestor first acquired the capacity of N_2_ fixation, it must have imposed a strong selection pressure on the individual cells. The trade-off between photosynthesis and nitrogen fixation led to the evolution of multiple solutions, which are still present in today’s cyanobacteria: the circadian rhythm of N_2_ fixation in unicellular cyanobacteria (Mitsui et al. 1986), specific cells devoted to N_2_ fixation in undifferentiated filaments (Berman-Frank et al. 2003; Bergman et al. 2013), and the differentiation of the highly specialized heterocyst in filamentous cyanobacteria (Flores et al. 2018).

The need to compartmentalize the two incompatible functions, photosynthesis and N_2_ fixation, has been proposed to drive the emergence of multicellular forms in cyanobacteria (Ispolatov et al. 2012). More specifically, the result from a quantitative theoretical model predicts that within a population of genetically identical unicellular nitrogen fixing cyanobacteria, cell differentiation and phenotypic heterogeneity would have been adaptive if this increased the fitness of the organisms in multicellular groups. In the case of unicellular cyanobacteria this means that cells evolved adhesion and exchanged fixed nitrogen and carbon products within early cell groups, such as filaments.

In filamentous cyanobacteria, dividing cells remain linked in a chain, resulting in a localization of cells in close spatial proximity, facilitating metabolite exchange between the individual cells. When compared to the more transient associations in spatially structured communities, such as in EPS imbedded biofilms, the development of filaments opens possibilities for a more direct and permanent exchange of molecules between neighboring cells with high specificity. Metabolic exchange could have evolved as described for the evolution of metabolic cross-feeding (D’Souza et al. 2018), as the exchange of carbon and nitrogen against other products is generally common in photosynthetic or nitrogen-fixing organisms (Kaiser et al. 2015).

The emergence of GAS VESICLES and MOTILITY traits signify the evolution from a stationary to a more active lifestyle, enabling cells to regulate their buoyancy in the water column. This result is further supported by the subsequent inference of UNATTACHED, which indicates the transition from a benthic to a planktonic lifestyle.

Thereafter the traits HORMOGONIA and EPIPHYTIC are inferred to occur simultaneously. The differentiation of HORMOGONIA can be induced by environmental stimuli, such as nitrogen deprivation (Flores & Herrero 2010). HORMOGONIA are released from the mother filament through the formation of necridia, dead cells resulting from PCD (Nürnberg et al. 2014). After their release from the main trichome, HORMOGONIA disperse via gliding motility or float thanks to GAS VESICLES, ensuring the reproduction of benthic species (Rippka et al. 1979).

HORMOGONIA with GAS VESICLES are thus important for distribution in aquatic environments as known from modern cyanobacteria (*Fischerella, Hapalosiphon, Tolypothrix*) (Komárek 2013). The close local association with plants, as indicated by EPIPHYTIC, however might have been the first step towards the initiation of one of the many symbioses between higher organisms and cyanobacteria, where HORMOGONIA serve as the infection units (Meeks & Elhai 2002).

Notably, the differentiation into HORMOGONIA is reversible, as they develop a sessile lifestyle, where they grow into a new vegetative filament (Flores & Herrero 2010). Here we observe the emergence of a life cycle with two distinct cell types, which is important for the transition to multicellularity (Hammerschmidt et al. 2014; Rose 2020). Such a life cycle results in selection operating at the higher, the filament level and thus ensures the reproduction of the newly formed collective entity.

### The evolution of cell differentiation leads to higher cyanobacterial complexity

A central innovation that is associated with this phase (iv) in the species tree is MULTIPLANE FISSION. This trait co-occurred with the ability to produce BAEOCYTES, differentiated cells, which are the reproductive stages in the order Pleurocapsales (Waterbury & Stanier 1978). Notably, baeocyte-forming cyanobacteria, that have been traditionally grouped together with unicellular cyanobacteria (Rippka et al. 1979), appear to immediately predate the evolution of spore-like AKINETES and nitrogen-fixing HETEROCYSTS and thus emerge much later than filamentous forms. Indeed, a recent study suggested that *Gloeocapsopsis* sp., a baeocytous cyanobacterium, harbors several characteristics that are in common with filamentous cyanobacteria, including mechanisms of cell-cell communication (Urrejola et al. 2020).

The late timing and the co-occurrence of the two traits AKINETES and HETEROCYSTS, indicative of higher complexity, are in line with the view that the evolution of the heterocyst was relatively late in the history of filamentous cyanobacteria (Tomitani et al. 2006), and where a common origin of akinetes and heterocysts has been proposed (Adams & Duggan 1999). HETEROCYSTS represent not only a morphological adaptation to the obstacle of N_2_ fixation under oxic conditions but also an elaborate and highly specialized communication and metabolite exchange system. In *Anabaena* sp., for example, where several hundred cells communicate within a filament, a regular heterocyst formation pattern along the filament must be achieved to guarantee that every cell is adequately supplied with fixed nitrogen compounds (Herrero et al. 2016). For this, the inhibitory signaling peptide PatS needs to be distributed along the filament with heterocyst formation occurring only in cells with low PatS concentration (Yoon & Golden 1998). Whether the exchange of metabolites and regulators happens via the continuous periplasm (Flores et al. 2006) or through septal junctions (Mullineaux et al. 2008) is still not fully resolved.

The trait that evolved last, based on the analysis, is TRUE-BRANCHING, where cells in a filament perform MULTIPLANE FISSION. TRUE-BRANCHING is characteristic for the members of the Haphalosiphon/Stigonematales clade. Our results confirm previous morphological and phylogenetic studies that found TRUE-BRANCHING to be the latest evolutionary innovation in cyanobacteria (Rippka et al. 1979; Dagan et al. 2013; Koch et al. 2017).

### Trade offs between incompatible processes lead to division of labor and stable multicellularity

Common features of evolutionary transitions in individuality comprise cooperation between the lower level units (Bonner 1998) and the division of labor (Michod 2007). The latter might be of particular advantage, and serve as the driver of the transition to multicellularity when there is a strong trade-off between processes that cannot be performed in a single cell at one time (Michod 2007; Ispolatov et al. 2012). Our current findings support this theory and point to nitrogen fixation, and its incompatibility with photosynthesis, as the trigger for the evolution of multicellularity in cyanobacteria. One open question concerns how the underlying genetics of novel traits, such as the division of labor, arise within a newly emerging multicellular individual. In the case of cyanobacteria multicellularity, as also suggested for animal multicellularity (Brunet & King 2017), we conjecture that no new genes were required and that higher complexity was achieved by regulatory changes in gene expression patterns. Basic communication and metabolite exchange was pre-existing as single-celled bacteria frequently engage in cell-cell communication and cross-feeding of metabolites via the external environment (D’Souza et al. 2018). Division of labor between photosynthesis and nitrogen fixation was likely first established by the regulatory mechanism of temporal switching. Once simple forms of division of labor and metabolic exchange existed, the transition into spatial separation in differentiated cells could have evolved mainly by regulatory modifications.

Differentiated cells are one of the hallmarks of complex multicellularity. It is therefore significant that we observe six distinct cell types in cyanobacteria: photosynthetic, hormogonia, necridia, akinetes, baeocytes, and heterocysts. Such a plurality indicates that the underlying regulatory mechanisms are well developed and that their plasticity and adaptability are a matter of course. It is also significant that three of the differentiated cell types, hormogonia, akinetes, and baeocytes, offer novel reproductive potential and the establishment of a multicellular life cycle. Moreover, signs of a nascent developmental plan can be observed in both the distribution of heterocysts along filaments and in the patterning of true branching cyanobacteria. These elements have no fitness value for the individual cell, but are selectable adaptations on the higher level, the filament. The chronology of the evolution of multicellularity in cyanobacteria shows that, once established, multicellular individuality opens new vistas of opportunities.

## Acknowledgements

We thank Tom Williams, Anne Kupczok, Tanita Wein, Claudia Taubenheim, Peter Deines, and Caroline Rose for comments on previous versions of the manuscript. We thank Fenna Stücker for illustrations. This work was supported by European Union’s Framework Programme for Research and Innovation Horizon 2020 (2014–2020) under the Marie Skłodowska-Curie Grant Agreement No. 657096, by the German Academic Exchange Service (DAAD) to KH, by CAPES (Coordination for the Improvement of Higher Education Personnel–Brazil) to F.D.K.T., and by the Chilean National Agency for Research and Development (ANID) Doctoral scholarship N° 21191763 to J.A..

## Author Contributions

K.H. and T.D. conceived the study. K.H. collected the traits data. G.L., F.D.K.T and J.A. performed the analyses. K.H., G.L., and T.D. wrote the manuscript with contributions from F.D.K.T. and J.A..

---

**Figure S1.**
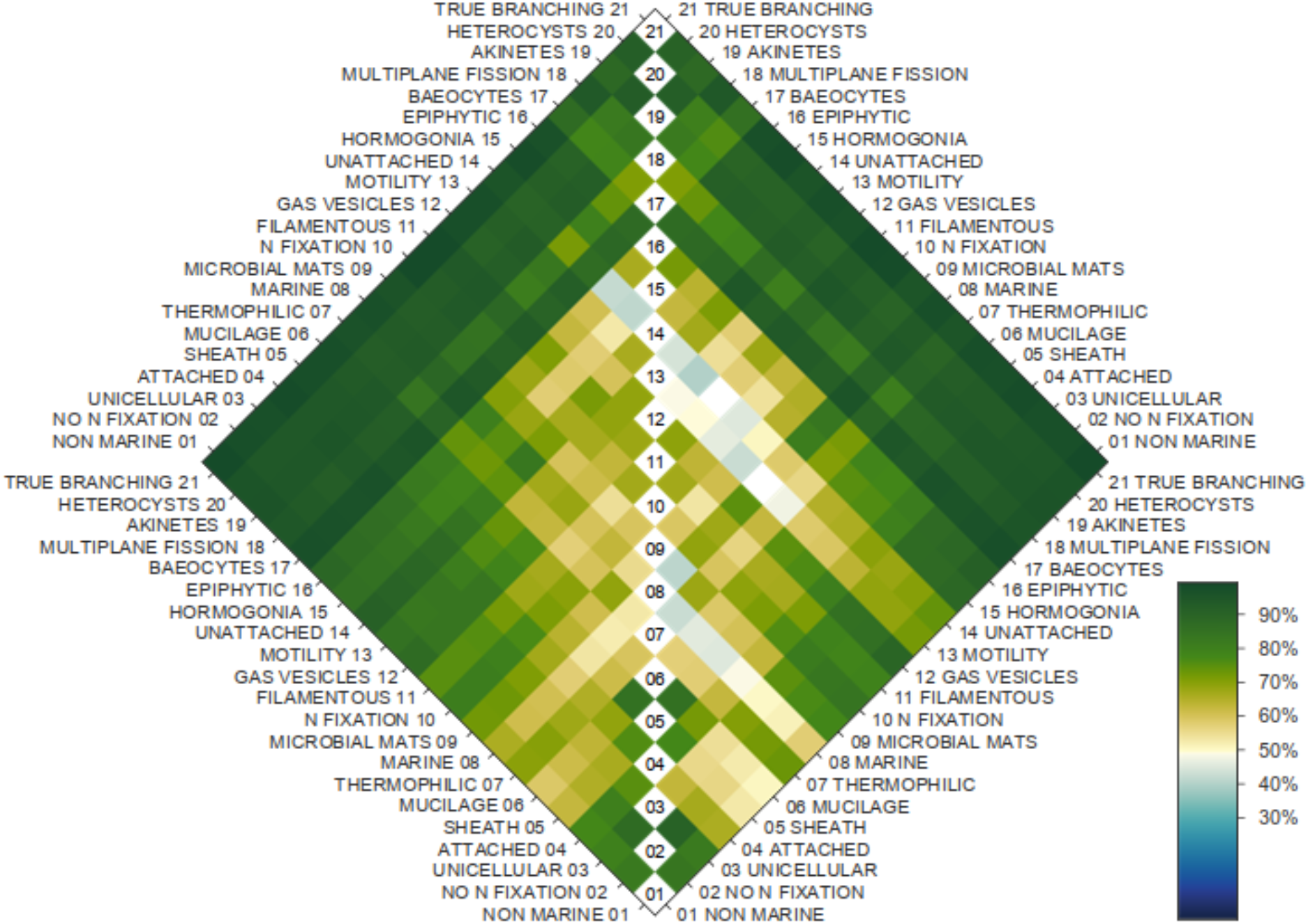
Gene tree support for the relative order of pairs of traits. Left: support from gene trees with SynPro clade excluded (167 spp.). Right: original gene trees as in fig. 1 (199 spp.). Cells in the matrix are shaded according to the proportion of gene trees that support the conclusion based on the species tree (according to the color bar on the right).

